# Multiple Long-read Sequencing Survey of Herpes Simplex Virus Lytic Transcriptome

**DOI:** 10.1101/605956

**Authors:** Dóra Tombácz, Zsolt Balázs, Gábor Gulyás, Zsolt Csabai, Miklós Boldogkoi, Michael Snyder, Zsolt Boldogkoi

**Author notes:** To whom correspondence should be addressed, Emails: DT, ZBa, GG, MB, ZC, MS, ZBo.

## Abstract

Long-read sequencing (LRS) has become increasingly important in RNA research due to its strength in resolving complex transcriptomic architectures. In this regard, currently two LRS platforms have demonstrated adequate performance: the Single Molecule Real-Time Sequencing by Pacific Biosciences (PacBio) and the nanopore sequencing by Oxford Nanopore Technologies (ONT). Even though these techniques produce lower coverage and are more error prone than short-read sequencing, they continue to be more successful in identifying transcript isoforms including polycistronic and multi-spliced RNA molecules, as well as transcript overlaps. Recent reports have successfully applied LRS for the investigation of the transcriptome of viruses belonging to various families. These studies have substantially increased the number of previously known viral RNA molecules. In this work, we used the Sequel and MinION technique from PacBio and ONT, respectively, to characterize the lytic transcriptome of the herpes simplex virus type 1 (HSV-1). In most samples, we analyzed the poly(A) fraction of the transcriptome, but we also performed random oligonucleotide-based sequencing. Besides cDNA sequencing, we also carried out native RNA sequencing. Our investigations identified more than 160 previously undetected transcripts, including coding and non-coding RNAs, multi-splice transcripts, as well as polycistronic and complex transcripts. Furthermore, we determined previously unsubstantiated transcriptional start sites, polyadenylation sites, and splice sites. A large number of novel transcriptional overlaps were also detected. Random-primed sequencing revealed that each convergent gene pair produces non-polyadenylated read-through RNAs overlapping the partner genes. Furthermore, we identified novel replication-associated transcripts overlapping the HSV-1 replication origins, and novel LAT variants with very long 5’ regions, which are co-terminal with the LAT-0.7kb transcript. Overall, our results demonstrated that the HSV-1 transcripts form an extremely complex pattern of overlaps, and that entire viral genome is transcriptionally active. In most viral genes, if not in all, both DNA strands are expressed.

## INTRODUCTION

Next-generation short-read sequencing (SRS) technology has revolutionized the research fields of genomics and transcriptomics due to its capacity of sequencing a large number of nucleic acid fragments simultaneously at a relatively low cost. Long-read sequencing (LRS) technology is able to read full-length RNA molecules, therefore it is ideal for application in the analysis of complex transcriptomic profiles. Currently two techniques are available in the market, the California-based Pacific Biosciences (PacBio) and the British Oxford Nanopore Technologies (ONT) platforms. The PacBio approach is based on single-molecule real-time (SMRT) technology, while the ONT platform utilizes the nanopore sequencing concept. Both techniques have already been applied for the transcriptomic analysis of various organisms (Byrne et al., 2017; Chen et al., 2017; Cheng et al., 2017; Li et al., 2018; Nudelman et al., 2018; Wen et al., 2018; Zhang et al., 2018), including viruses, such as herpesviruses (Balázs et al., 2017a, 2017b; Depledge et al., 2018; Moldován et al., 2017b; O’Grady et al., 2016; Tombácz et al., 2015, 2016, 2017b, 2017a, 2018d), poxviruses (Tombácz et al., 2018c), baculoviruses (Moldován et al., 2018b), retroviruses (Moldován et al., 2018a) and circoviruses (Moldován et al., 2017a).

Herpes simplex virus type 1 (HSV-1) is a human pathogenic virus belonging to the *Alphaherpesvirinae* subfamily of the *Herpesviridae* family. Its closest relatives are the HSV-2, the Varicella-zoster virus (VZV), and the animal pathogen pseudorabies virus (PRV). The most common symptom of HSV-1 infection is cold sores, which can recur from latency causing blisters primarily on the lips. HSV-1 may cause acute encephalitis in immunocompromised patients. The ability of herpesviruses to establish lifelong latency within the host organism significantly contributes to their evolutionary success: according to WHO’s estimates, more than 3.7 billion people under the age of fifty are infected with HSV-1 worldwide (Looker et al., 2015).

HSV-1 has a 152-kbp linear double-stranded DNA genome that is composed of unique and repeat regions. Both the long (UL) and the short (US) unique regions are flanked by inverted repeats (IRLs and IRSs) (Macdonald et al., 2012). The viral genome is transcribed by the host RNA polymerase in a cascade-like manner producing three kinetic classes of transcripts and proteins: immediate-early (IE), early (E) and late (L) (Harkness et al., 2014). The IE genes encode transcription factors required for the expression of E and L genes. The E genes mainly code for proteins playing a role in the DNA synthesis, whereas the L genes specify the structural elements of the virus. Earlier studies and *in silico* annotations have revealed 89 mRNAs, 10 non-coding (nc)RNAs (Hu et al., 2016; Lim, 2013; Macdonald et al., 2012; McGeoch et al., 2006; Rajčáni et al., 2004), and 18 microRNAs (Du et al., 2015). Our recent study (Tombácz et al., 2017b) based on PacBio RS II sequencing has identified additional 142 transcripts and transcript isoforms, including ncRNAs. The detection and the kinetic characterization of HSV-1 transcriptome face an important challenge because of the overlapping and polycistronic nature of the viral transcripts. The polycistronic transcription units are different from those of the bacterial operons, in that the downstream genes on the multigenic transcripts are untranslated because herpesvirus mRNAs use cap-dependent translation initiation (Merrick, 2004). The majority of herpesvirus transcripts are organized into tandem gene clusters generating overlapping transcripts with co-terminal transcription end sites (TESs). The *ul40-44* genomic region of HSV-1 does not follow this rule, since these genes are primarily expressed as monocistronic RNA molecules. Our earlier study has revealed that these genes also produce low-abundance bi- and polycistronic transcripts. Alternatively, many HSV-1 genes, which were believed to be exclusively expressed as parts of multigenic RNAs, have been shown to also specify low-abundance monocistronic transcripts (Tombácz et al., 2017b).

SRS technologies have become useful tools for the analysis of the transcriptomes. However, the conventionally applied SRS platforms cannot reliably distinguish between multi-spliced transcript isoforms, and the variants of the transcription start sites (TSS), as well as between the embedded transcripts and their host RNAs, etc. Additionally, SRS, even if applied in conjunction with auxiliary techniques such as RACE analysis, has limitations in detecting multigenic transcripts, including polycistronic RNAs and complex transcripts (cxRNAs; containing genes standing in opposite orientations). LRS is able to circumvent these problems. Both PacBio and ONT approaches are capable of reading cDNAs generated from full-length transcripts in a single sequencing run and permit mapping of TSSs and TESs with base-pair precision. The most important disadvantage of LRS compared to SRS techniques is the lower coverage. In PacBio sequencing, if any errors occur in the raw reads, they are easily corrected thanks to the very high consensus accuracy of this technique (Miyamoto et al., 2014). Thus, it is only a widespread myth that SMRT sequencing is too error prone to be used for precise sequence analysis. The precision of basecalling is substantially lower for the ONT platform than that of the PacBio, but the former technique is far more cost-effective, and yields both higher throughput and longer reads. The high error rate of ONT technique can be circumvented by obtaining high sequence coverage. Nonetheless, this latter problem is not a critical in transcriptome research if the genome sequence of the examined organism has already been annotated.

A diverse collection of methods and approaches has already been employed for the investigation of the herpesvirus transcriptomes, including *in silico* detection of open reading frames (ORFs) and cis-regulatory motifs, Northern-blot analysis (Costa et al., 1984; Sedlackova et al., 2008), S1 nuclease mapping (McKnight, 1980; Rixon and Clements, 1982), primer extension (Naito et al., 2005; Perng et al., 2002), real-time reverse transcription-PCR (RT^2^-PCR) analysis (Tombácz et al., 2009), microarrays (Stingley et al., 2000), Illumina sequencing (Harkness et al., 2014; Oláh et al., 2015), PacBio RS II (O’Grady et al., 2016; Tombácz et al., 2016, 2017b) and Sequel sequencing, as well as ONT MinION sequencing (Boldogkői et al., 2018b; Prazsák et al., 2018).

In this study, we report the application of PacBio Sequel and ONT MinION long-read sequencing technologies for the characterization of the HSV-1 lytic transcriptome. We used an amplified isoform sequencing (Iso-Seq) protocol of PacBio that was based on PCR amplification of the cDNAs prior to sequencing. We used both cDNA and direct (d)RNA sequencing of the ONT platform. Additionally, we applied Cap-selection for ONT sequencing. In order to identify non-polyadenylated transcripts, we also applied random oligonucleotide primer-based RT in addition to the oligo(dT)-priming. Furthermore, this latter technique is more efficient for mapping of the TSSs, and it is useful for the validation of novel RNA molecules. Our intention of using novel LRS techniques was to generate a higher number of sequencing reads as well as to identify novel transcripts that had been undetected in our earlier PacBio RS II-based approach. Furthermore, in this report, we reanalyzed our earlier results that we had obtained by a single-platform method (Tombácz et al., 2017b).

## MATERIALS AND METHODS

### Cells and viral infection

The strain KOS of HSV-1 was propagated on an immortalized kidney epithelial cell line (Vero) isolated from the African green monkey (*Chlorocebus sabaeus*). Vero cells were cultivated in Dulbecco’s modified Eagle medium supplemented with 10% fetal bovine serum (Gibco Invitrogen) and 100μl penicillin-streptomycin 10K/10K mixture (Lonza) /ml and 5% CO_2_ at 37°C. The viral stocks were prepared by infecting rapidly-growing semi-confluent Vero cells at a multiplicity of infection (MOI) of 1 plaque-forming unit (pfu)/cell, followed by incubation until a complete cytopathic effect was observed. The infected cells were then frozen and thawed three times, followed by spinning at 10,000xg for 15min using low-speed centrifugation. For the sequencing studies cells were infected with MOI=1, incubated for 1h, followed by removal of the virus suspension and washing with PBS. This was followed by the addition of fresh culture medium to the cells. The cells were incubated for 1, 2, 4, 6, 8, 10, 12 or 24h.

### RNA isolation

The total RNA samples were purified from the cells using the NucleoSpin® RNA kit (**Table 1**) according to the kit’s manual and our previously described methods (Boldogkői et al., 2018a; Tombácz et al., 2018d, 2018c). The RNA samples were quantified by using the Qubit® 2.0 Fluorometer and they were stored at −80°C until use. The samples taken from each experiment were then mixed for sequencing. Samples were subjected to ribodepletion for the random primed sequencing, while selection of the poly(A)^+^ RNA fraction was carried out for polyA-sequencing. All experiments were performed in accordance with relevant guidelines and regulations.

**Table 1.** Summary of the kits used for RNA preparation and quantitation.

|  | Method | Kit | Company |
| --- | --- | --- | --- |
| RNA purification | Total RNA extraction | NucleoSpin RNA | Macherey Nagel |
|  | PolyA(+) RNA isolation | Oligotex mRNA Mini Kit | Qiagen |
|  | Ribodepletion | Ribo-Zero™ Magnetic Kit H/M/R | Epicentre/Illumina |
| Concentration measurement | Total RNA | Qubit RNA BR Assay Kit | Life Technologies |
|  | PolyA(+) RNA | Qubit RNA HS Assay Kit |  |
|  | rRNA depleted RNA |  |  |

### Pacific RS II and Sequel platforms – sequencing of the polyadenylated RNA fraction or the total transcriptome

The Clontech SMARTer PCR cDNA Synthesis Kit was used for cDNA preparation according to the PacBio Isoform Sequencing (Iso-Seq) protocol. For analysis of the relatively short viral RNAs, the ‘No-size selection’ method was followed and samples were run on both the RSII and Sequel platforms. The SageELF™ and BluePippin™ Size-Selection Systems (Sage Science) were also used to carry out size-selection for capturing the potential long, rare transcripts. The reverse transcription (RT) reactions were primed by using the oligo(dT) from the SMARTer Kit, however we also used random primers for a non-size selected sample for the detection of non-polyadenylated RNAs. The cDNAs were amplified by using the KAPA HiFi Enzyme from KAPA Biosystems, according to the PacBio’s recommendations (Balázs et al., 2017b; Tombácz et al., 2018d). The SMRTbell libraries were generated with the PacBio Template Prep Kit 1.0. For binding the DNA polymerase and annealing the sequencing primers, the DNA/Polymerase Binding Kit P6-C4 and v2 primers, as well as the Sequel Sequencing Kit and v3 primers were used for the RSII and Sequel sequencing, respectively. The polymerase-template complexes were bound to MagBeads with the PacBio MagBead Binding Kit. Samples were loaded onto the RSII SMRT Cell 8Pac v3 or Sequel SMRT Cell 1M. The movie time was 240 or 360 minutes per SMRT Cell for the RSII, while 600 minutes movie time was set to the Sequel run.

### Oxford Nanopore MinION platform – sequencing of cDNA using oligo(dT) or random primers

#### Regular (no cap selection) protocol

The 1D Strand switching cDNA by ligation protocol (Version: SSE_9011_v108_revS_18Oct2016) from the ONT was used for sequencing the HSV-1 cDNAs on the MinION platform. The ONT Ligation Sequencing Kit 1D (SQK-LSK108) was applied for the library preparation using the recommended oligo(dT) primers, or custom made random oligonucleotides, as well as SuperScipt IV enzyme for the RTs. The cDNA samples were subjected to PCR reaction with KAPA HiFi DNA Polymerase (Kapa Biosystems) and Ligation Sequencing Kit Primer Mix (part of the 1D Kit). NEBNext End repair/dA-tailing Module (New England Biolabs) was used for the end repair, whereas the NEB Blunt/TA Ligase Master Mix (New England Biolabs) was utilized for the adapter ligation. The enzymatic steps (e. g.: RT, PCR, ligation) were carried out in a Veriti Cycler (Applied Biosystems) according to the 1D protocol (Moldován et al., 2018b; Tombácz et al., 2018d). The Agencourt AMPureXP system (Beckman Coulter) was used for purification of the samples after each enzymatic reaction. The quantity of the libraries was checked using the Qubit Fluorometer 2.0 and the Qubit (ds)DNAHS Assay Kit (Life Technologies). The samples were run on the R9.4 SpotON Flow Cells from ONT.

#### Cap selection protocol

The TeloPrime Full-Length cDNA Amplification Kit (Lexogen) was used for generating cDNAs from 5’ capped polyA^(+)^ RNAs. The RT reactions were carried out with oligo(dT) primers (from the kit) or random hexamers (custom made) using the enzyme from the kit. A specific adapter (capturing the 5’ cap structure) was ligated to the cDNAs (25°C, overnight), then the samples were amplified by PCR using the Enzyme Mix and the Second-Strand Mix from the TeloPrime Kit. The reactions were performed in a Veriti Cycler and the samples were purified on silica membranes (TeloPrime Kit) after the enzymatic reactions. The Qubit 2.0 and the Qubit dsDNA HS quantitation assay (Life Technologies) were used for measuring the concentration of the samples. A quantitative PCR reaction was carried out for checking the specificity of the samples using the Rotor-Gene Q cycler (Qiagen) and the ABsolute qPCR SYBR Green Mix from Thermo Fisher Scientific. A gene-specific primer pair (HSV-1 *us9* gene, custom made) was used for the test amplification. The PCR products were used for ONT library preparation and sequencing. The end-repair and adapter ligation steps were carried out as was described in the ‘Regular’ protocol, and in our earlier publication (Tombácz et al., 2018d, 2018c). The ONT R9.4 SpotONFlow Cells were used for sequencing.

### Oxford Nanopore MinION platform - Direct RNA sequencing

The ONT’s Direct RNA sequencing (DRS) protocol (Version: DRS_9026_v1_revM_15Dec2016) was used to examine the transcript isoforms without enzymatic reactions - to avoid the potential biases - as well as to identify possible base modifications alongside the nucleotide sequences. Polyadenylated RNA was extracted from the total RNA samples and it was subjected to DRS library preparation according to the ONT’s protocol (Tombácz et al., 2018d). The quantity of the sample was measured by Qubit 2.0 Fluorometer using the Qubit dsDNA HS Assay Kit (both from Life Technologies). The library was run on an ONT R9.4 SpotON Flow Cell.

### Mapping and data analysis

The minimap2 aligner (Li, 2018) was used with options *-ax splice -Y -C5 --cs* for mapping the raw reads to the reference genome (X14112.1), followed by the application of LoRTIA toolkit (https://github.com/zsolt-balazs/LoRTIA) for the determination of introns, the 5’ and 3’ ends of the transcripts, as well as for detecting the full-length reads. Introns were defined as deletions with the consensus flanking sequences (GT/AG, GC/AG, AT/AC). The complete intron lists are available as additional material (**Supplementary Table 1**). We used even more strict criteria, those splice sites were accepted, which were validated by dRNA-Seq [used in our present work and in Depledge and coworkers’ study (Depledge et al., 2018]: these transcripts all have the canonical splice site: GT/AG and they are abundant (>100 read in Sequel data).

The 5’ adapter and the poly(A) tail sequences were identified at the ends of the reads by the LoRTIA toolkit based on the Smith-Waterman alignment scores (**Table 2**). If the adapter or poly(A) sequence ended at least 3 nucleotides (nts) downstream from the start of the alignment, the adapter was discarded, presuming that it could have been placed there by template-switching. Transcript features such as introns, transcriptional start sites (TSS) and transcriptional end sites (TES) were annotated if they were detected in at least two reads and 0.1% of the local coverage. In order to reduce the effects of RNA degradation, only those TSSs were annotated, which were significant peaks compared to their ±50-nt-long windows according to a Poisson distribution. Reads being connected a unique set of transcript features were annotated as transcript isoforms. Low-abundance reads detected in a single experiment were accepted as transcripts if the same TSS and TES were also used by other transcripts. In most cases those reads were accepted as isoforms, which were detected in at least two independent experiments. The 5′-ends of the long low-abundance reads were checked individually using the Integrative Genome Viewer (IGV; https://software.broadinstitute.org/software/igv/download).

**Table 2.**
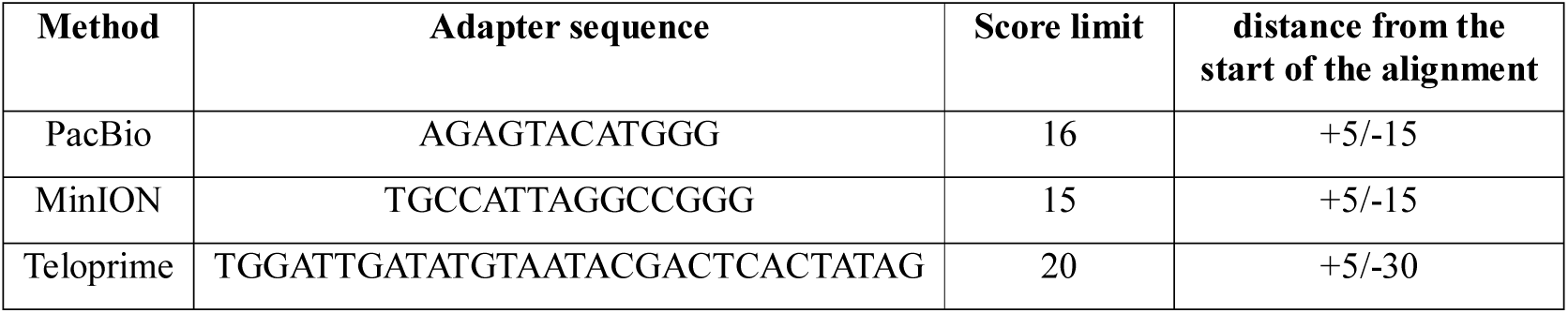
The 5’ adapter sequences and settings for adapter detection with the LoRTIA pipeline. The scoring of the Smith-Waterman alignment was set to +2 for matches and −3 for mismatches, gap opening and gap extensions.

## RESULTS

### Analysis of the HSV-1 transcriptome with full-length sequencing

In this work, we report the application of two LRS techniques, the PacBio Sequel and the ONT MinION platforms, for the investigation of the HSV-1 lytic transcriptome. We also reutilized our previous PacBio RS II data for the validation of the novel transcripts. The PacBio sequencing is based on amplified Iso-Seq template preparation protocol that utilizes a switching mechanism at the 5’ end of the RNA template, thereby producing complete full-length cDNAs (Zhu et al., 2001). We applied both cDNA and dRNA sequencing for the ONT technique. Additionally, we used Cap-selection for a fraction of samples. The sequencing reads were mapped to the HSV-1 (X14112) genome using the Minimap2 alignment tool (Li, 2018) with default parameters.

Altogether, we obtained 67,688 full-length ROIs mapped to the HSV-1 genome by the Sequel sequencing with a mean length of 1,805 nts, whereas the PacBio RSII platform generated 31,623 ROIs with an average of 1,368 nts. The total number of ROIs aligned to the HSV-1 genome is 39,096 and 77,851 from the RSII and Sequel sequencing, respectively. **Table 3** shows the read lengths of full-length ROIs, as well as the average read lengths of the given samples.

**Table 3.** Average read-length obtained from the methods used in this study.

| Technique | Average length of the reads (bp) | Average length of the abundant full length transcripts (bp) |
| --- | --- | --- |
| PacBio RSII *oligo(dT)* | 1,536 | 1,368 |
| PacBio RSII *random* | 1,000 | NA |
| PacBio Sequel | 2,104 | 1,805 |
| ONT MinION 1D *oligo(dT)* | 1,466 | 1,259 |
| ONT MinION 1D *random* | 910 | NA |
| ONT MinION Cap-seq *oligo(dT)* | 1,067 | 810 |
| ONT MinION dRNA-Seq | 1,111 | NA |

ONT sequencing produced altogether 175,169 sequencing reads, however, the number of complete, full-length reads is much fewer (5,958 reads). The reason behind the relatively low full-length read count within the MinION samples is that this method – compared to the PacBio -generates a higher number of 5’ truncated reads. We have reported in our previous publications that the dRNA-Seq method is not suitable for capturing the entire transcripts (Moldován et al., 2017b, 2018b): we found that short 5’ sequences of the transcripts and the polyA-tails were missing from most of the reads.

Another drawback of native RNA sequencing is its low throughput compared to cDNA sequencing. The advantages of dRNA-Seq are that it is free of the false products typical to those produced by RT, PCR, and cDNA sequencing. Cap-selection did not perform very well in our experiment, because it produced relatively short average sequencing reads, which was similar to the results obtained in other herpesviruses [PRV (Tombácz et al., 2018d) and VZV (Tombácz et al., 2018b)]. However, this phenomenon has not been observed in baculoviruses (Moldován et al., 2018b) and vaccinia virus (Tombácz et al., 2018c). Random RT-priming allowed the analysis of non-polyadenylated transcripts, and helped the validation of the TSSs and the splice sites. Additionally, this technique proved to be superior for identifying the 5’-ends of very long transcripts, including polycistronic and complex RNA molecules.

The following technical artifacts can be generated by RT and PCR: template switching, as well as the non-specific binding of the oligod(T) or PCR primers. In addition to the poly(A) tails, the oligo(dT) primers occasionally hybridize to A-rich regions of the transcripts and thereby produce false reads. These products were discarded from further analysis, albeit in some cases we were unsure about the nonspecificity of the removed reads. We ran altogether 11 parallel sequencing reactions using eight different sequencing techniques for providing independent reads. Additionally, in some cases, the same TSS, TES or splice junction were found in other transcripts detected within the same sequencing reaction, which further enhanced the number of independent sequencing reads. In our earlier publication, we could not detect all of the spurious products, therefore, in our present work, we have made a minor correction to our formerly published results.

### Novel 5’-truncated mRNAs

Present investigations revealed 28 novel truncated mRNAs (tmRNAs) of HSV-1 (**Table 4**), which were all produced from genes embedded in larger host genes of the virus. These 5’-truncated mRNAs are generated by alternative transcription initiation from promoters located within the larger genes and contain in-frame ORFs. The first in-frame AUG triplets are assumed to encode the translation start codons. Further analyses have to be carried out to verify the coding potential of the ORF-containing tmRNAs.

**Table 4.**
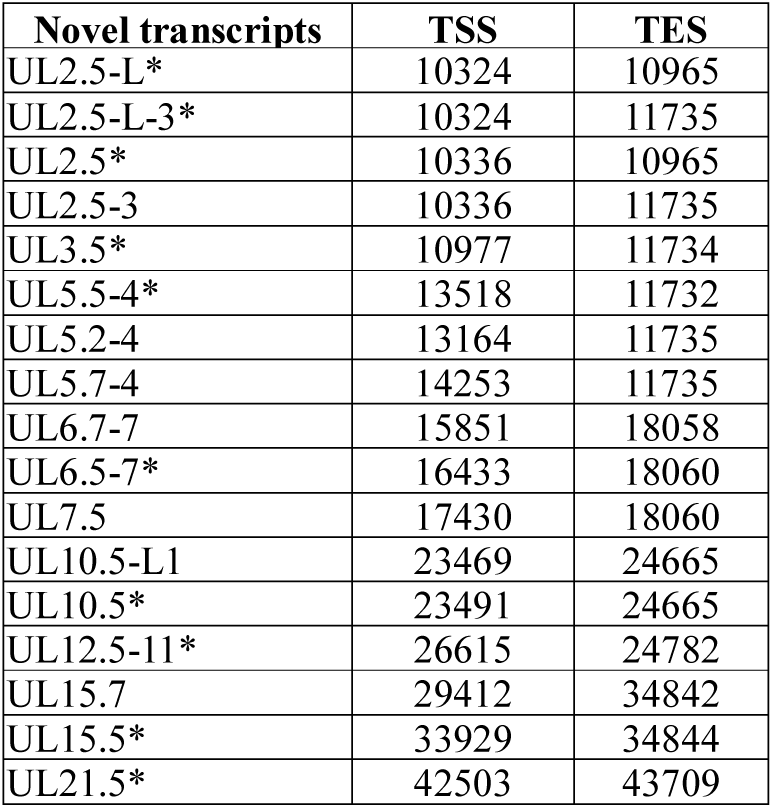
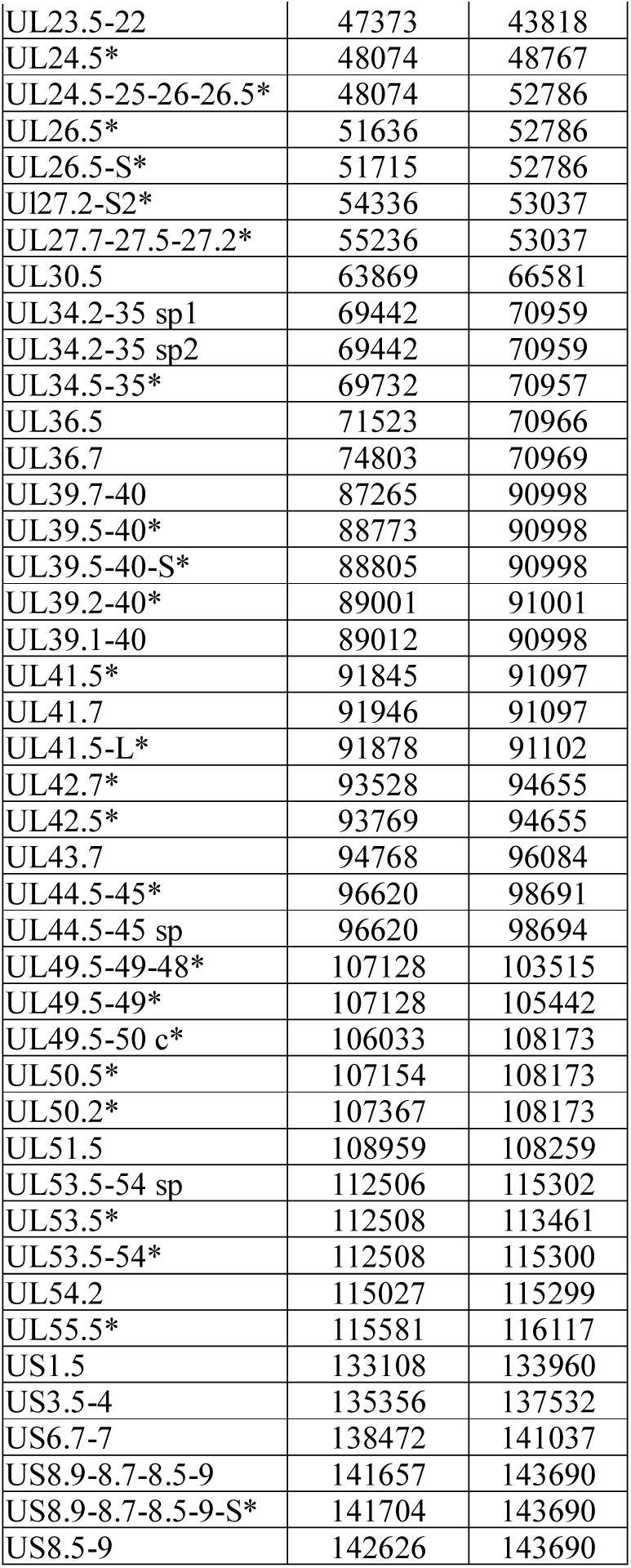
Novel HSV-1 transcripts with putative coding potential. This table summarizes the novel and the previously published embedded mRNAs, as well as their genomic positions. The asterisks indicate the detection of the given transcripts in our earlier study (Tombácz et al., 2017b).

| Novel transcripts | TSS | TES |
| --- | --- | --- |
| UL2.5-L* | 10324 | 10965 |
| UL2.5-L-3* | 10324 | 11735 |
| UL2.5* | 10336 | 10965 |
| UL2.5-3 | 10336 | 11735 |
| UL3.5* | 10977 | 11734 |
| UL5.5-4* | 13518 | 11732 |
| UL5.2-4 | 13164 | 11735 |
| UL5.7-4 | 14253 | 11735 |
| UL6.7-7 | 15851 | 18058 |
| UL6.5-7* | 16433 | 18060 |
| UL7.5 | 17430 | 18060 |
| UL10.5-L1 | 23469 | 24665 |
| UL10.5* | 23491 | 24665 |
| UL12.5-11* | 26615 | 24782 |
| UL15.7 | 29412 | 34842 |
| UL15.5* | 33929 | 34844 |
| UL21.5* | 42503 | 43709 |
| UL23.5-22 | 47373 | 43818 |
| UL24.5* | 48074 | 48767 |
| UL24.5-25-26-26.5* | 48074 | 52786 |
| UL26.5* | 51636 | 52786 |
| UL26.5-S* | 51715 | 52786 |
| UL27.2-S2* | 54336 | 53037 |
| UL27.7-27.5-27.2* | 55236 | 53037 |
| UL30.5 | 63869 | 66581 |
| UL34.2-35 sp1 | 69442 | 70959 |
| UL34.2-35 sp2 | 69442 | 70959 |
| UL34.5-35* | 69732 | 70957 |
| UL36.5 | 71523 | 70966 |
| UL36.7 | 74803 | 70969 |
| UL39.7-40 | 87265 | 90998 |
| UL39.5-40* | 88773 | 90998 |
| UL39.5-40-S* | 88805 | 90998 |
| UL39.2-40* | 89001 | 91001 |
| UL39.1-40 | 89012 | 90998 |
| UL41.5* | 91845 | 91097 |
| UL41.7 | 91946 | 91097 |
| UL41.5-L* | 91878 | 91102 |
| UL42.7* | 93528 | 94655 |
| UL42.5* | 93769 | 94655 |
| UL43.7 | 94768 | 96084 |
| UL44.5-45* | 96620 | 98691 |
| UL44.5-45 sp | 96620 | 98694 |
| UL49.5-49-48* | 107128 | 103515 |
| UL49.5-49* | 107128 | 105442 |
| UL49.5-50 c* | 106033 | 108173 |
| UL50.5* | 107154 | 108173 |
| UL50.2* | 107367 | 108173 |
| UL51.5 | 108959 | 108259 |
| UL53.5-54 sp | 112506 | 115302 |
| UL53.5* | 112508 | 113461 |
| UL53.5-54* | 112508 | 115300 |
| UL54.2 | 115027 | 115299 |
| UL55.5* | 115581 | 116117 |
| US1.5 | 133108 | 133960 |
| US3.5-4 | 135356 | 137532 |
| US6.7-7 | 138472 | 141037 |
| US8.9-8.7-8.5-9 | 141657 | 143690 |
| US8.9-8.7-8.5-9-S* | 141704 | 143690 |
| US8.5-9 | 142626 | 143690 |

### Novel non-coding transcripts

In this part of our study, we detected twelve non-coding RNAs, including antisense RNAs (asRNAs, labeled as ASTs) and other long non-coding RNAs (lncRNAs) (**Table 5**). Furthermore, we validated and determined the base pair-precision termini of the transcripts published earlier by us or by others.

**Table 5.**
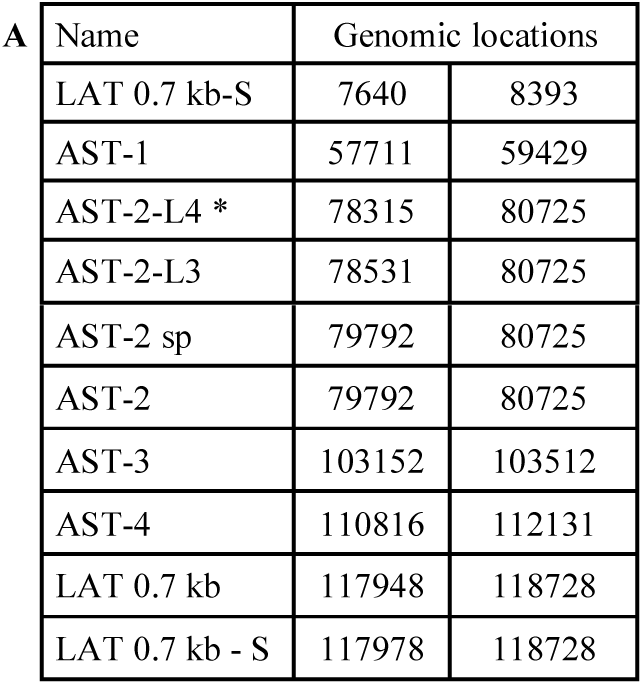
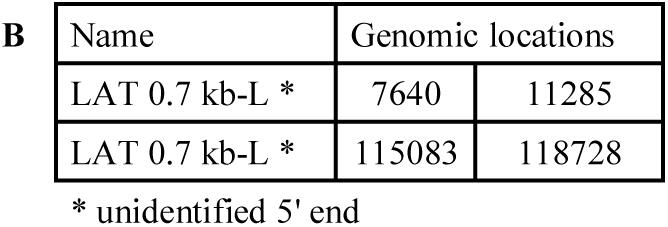
Polyadenylated ncRNAs of HSV-1. **A**: Previously detected and validated ncRNAs; **B**: Novel ncRNAs. All transcripts are polyadenylated.

#### (1) Antisense RNAs

These transcripts can be controlled by their own promoters or by a promoter of other genes. It has earlier been reported that the 0.7-kb LAT transcript is not expressed in strain KOS of HSV-1 (Zhu 1999). Here we demonstrate that this is not the case, since we could detect this transcript. The existence of the shorter LAT-0.7kb-S (Tombácz et al., 2017b) was also confirmed. Additionally, we detected asRNAs being co-terminal with the LAT-0.7 transcripts, but having much longer TSSs. The LAT region and the surrounding genomic sequences are illustrated in **Figure 1**. Using random oligonucleotide-based LRS, we obtained a large number of antisense-oriented reads, most of them without identified 5’-ends. We also detected antisense transcripts without determined TSS and TES within the following 24 HSV-1 genes (*rl1, rl2, ul1, ul2, ul4, ul5, ul10, ul14, ul15, ul19, ul23, ul29, ul32, ul36, ul37, ul39, ul42, ul43, ul44, ul49, ul53, ul54, us4, us5*). The expression level of these asRNAs is low, in most cases only a few reads were detected per gene locus. However, high antisense expression was identified within the locus of *ul10* gene. A special class of asRNAs includes overlapping RNAs produced by divergent genes, and read-through RNAs (rtRNAs) generated by transcriptional read-through between convergent gene pairs.

**Figure 1.**
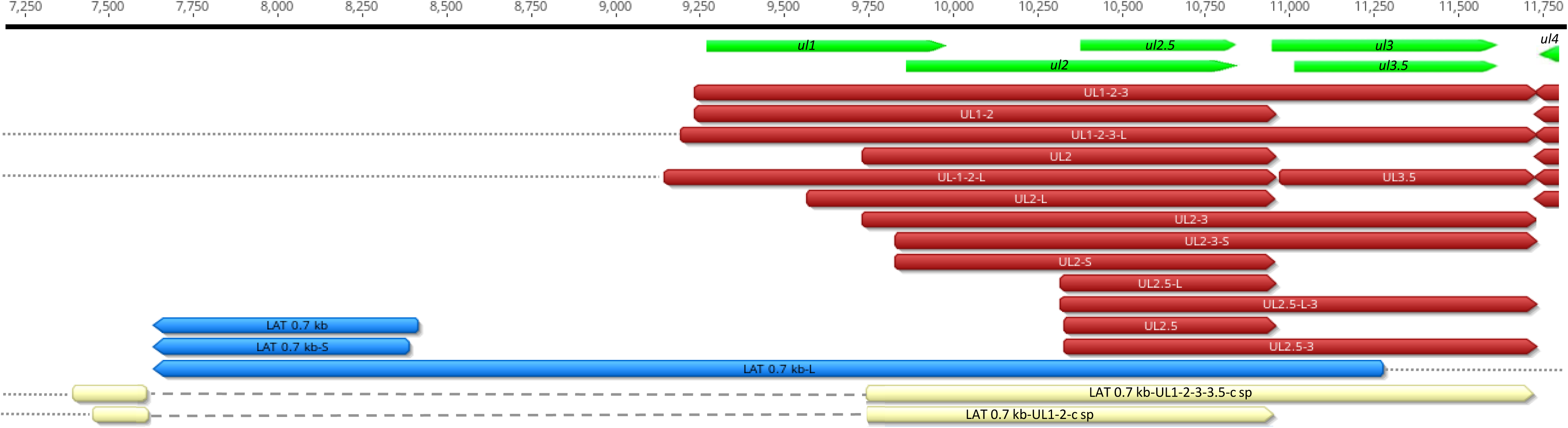
Schematic representation of the LAT region and surroundings. Besides the previously published coding and non–coding transcripts, this figure illustrates the newly discovered longer TSS version of the 0.7 kb LAT, as well as the oppositely oriented complex transcripts, which are co-terminal with the 3’ ends of the UL2 or UL3 transcripts. This transcriptional complexity is typical throughout the entire viral genome. Dashed lines: intron; dotted lines: unidentified TSS. The “c” tag stands for “complex”; “sp” for “spliced”; the “L” for “long”; and the “s” for short transcripts. The transcripts labeled by asterisks have unidentified TSSs, and thus they may be cxRNAs. The dotted lines represent introns, whereas the dashed lines indicate unidentified segments of the transcripts. Color legend: green - ORF; red - mRNA; blue - non-coding RNA; off-white - complex transcript

#### (2) Intergenic ncRNAs

A ncRNA (termed “unique long terminal ncRNA”; ULTN) with unidentified transcript ends was detected to be expressed from the outer termini of the HSV-1 unique long region. ULTN does not overlap any protein-coding genes. Their potential function remains unclear. A bidirectional, low-level expression from the intergenic region between the *rl2* (icp0) and LAT genes was also observed. These RNA molecules are co-terminal with the LAT-0.7kb transcript and may be parts of the potential RL2-LAT-UL1-2-3 transcript (Tombácz et al., 2017b). Additionally, we detected RNA expression in practically every intergenic region (**Table 5**). The *ul43-44-45/ul48-47- 46* convergent cluster is the only exception where we did detect neither read-through nor random overlapping RNAs.

#### (3) Replication-associated transcripts

We identified five replication-associated RNA (raRNA) designated OriL-RNA1, OriL-RNA2 and OriS-RNA1, which overlaps the replications origins OriL or OriS. OriL-RNA1 is a long TSS isoform produced from the *ul30* gene, whereas OriS-RNA2 is an *rs1* (*icp4*) TSS variant (**Figure 2**). OriL-RNA2 is a transcript without annotated TES. We suppose that this transcript is the long TSS variant of the *ul29* gene. We were able to detect only certain segments but not the entire OriS-RNA1 described by Voss and Roizman (Voss and Roizman, 1988). We also detected a longer TSS transcript isoform of the *us1* gene (US1-L2 = OriS-RNA3) which overlaps the OriS located within TRS (**Figure 2**). The OriS is also overlapped by a longer 5’ variant of the *us12* gene (US12-11-10-L2 = OriS-RNA4).

**Figure 2.**
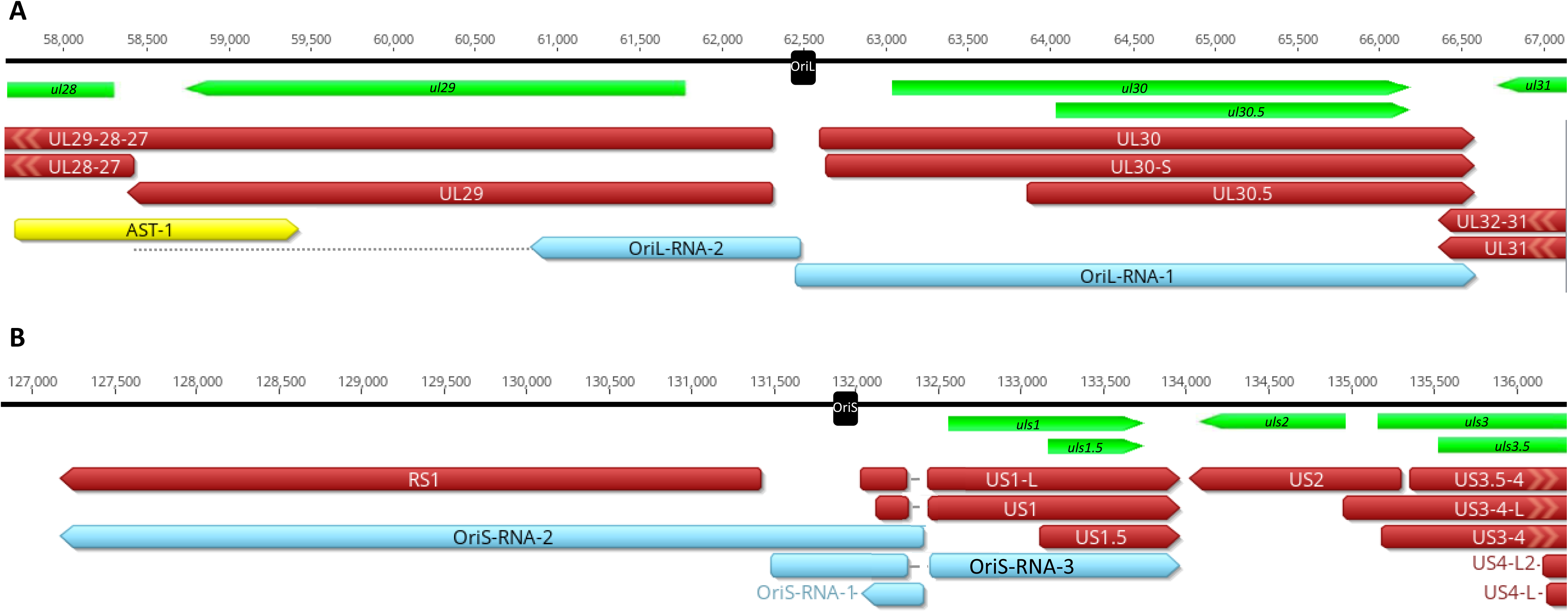
Replication associated transcripts of HSV. Color legend: green - ORF; red - mRNA; blue - non-coding RNA; yellow - antisense transcript; black rectangle: replication origin

### TSS and TES isoforms

The multiplatform system allowed the discovery of novel RNA isoforms and reannotation of the transcript termini published earlier by others and us (Depledge et al., 2018; Tombácz et al., 2017b). We used a novel bioinformatics tool (LoRTIA), developed in our laboratory, for the detection of TSS and TES positions. With this, we identified 218 TSS and 56 TES positions (**Supplementary Table 2 and 3**). Altogether 53 genes produce at least one TSS isoforms, besides the most frequent variants (**Supplementary Table 4**). Fifteen genes were found to produce three different transcript length isoforms (including the most frequent versions).

The recent LRS analysis discovered 51 protein-coding and two (0.7 kb LAT, and RS1) non-coding transcripts with alternative TSSs. However, a few transcripts with unannotated 5’-ends were also detected (**Supplementary Table 4**). The alternative TSSs may lead to transcriptional overlap or enlarge the extent of existing overlaps especially between divergently transcribed genes (**Figure 3**). The *ul21* gene produces nine different 5’ length variants, the longer ones overlap the divergently oriented *ul22* gene). Additionally, long TSS isoforms are responsible for the overlaps of each replication origin. The UL21 and UL10 transcripts exhibit an especially high complexity of TSS isoforms (**Figure 3**). Many of the longer TSS variants contain upstream ORFs (uORFs), which may carry distinct coding potentials as described by Balázs and colleagues in human cytomegalovirus (Balázs et al., 2017a). Two novel 3’-UTR variants were also identified in this study.

**Figure 3.**
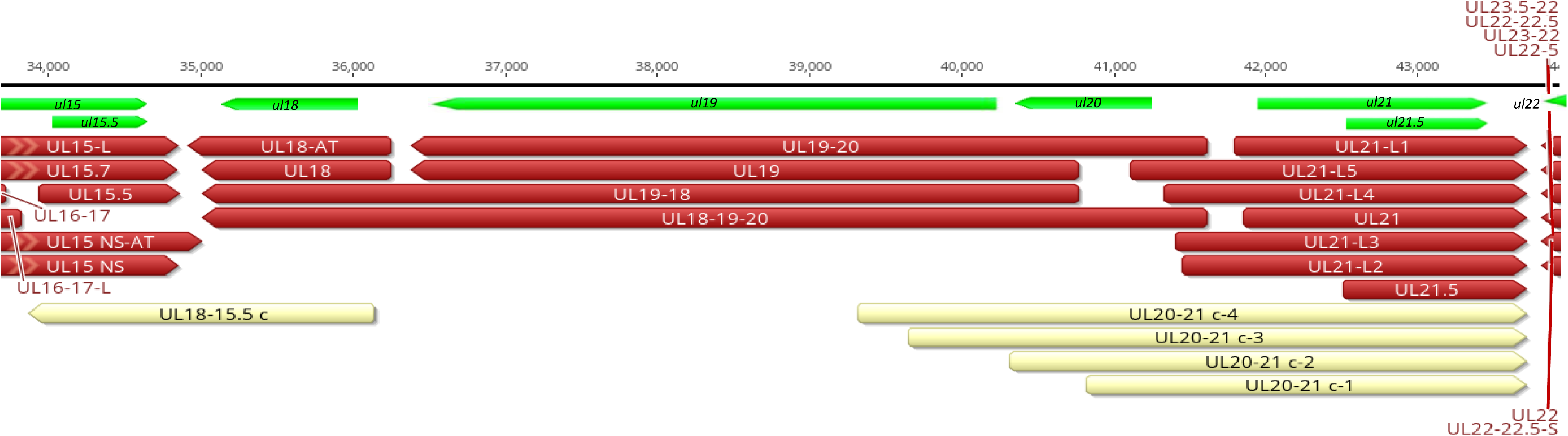
Divergent overlaps between the ul20 and ul21 genes. Color legend: green - ORF; red - mRNA; off-white - complex transcript

### Novel splice sites and splice isoforms

In this study, we also used dRNA sequencing, which does not produce spurious splice sites. The splice sites were confirmed by using the LoRTIA tool. Altogether, using the different sequencing techniques and bioinformatics analysis, we were able to verify the existence of nine previously described and four novel splice sites. These splice sites listed in **Table 6** are frequent and most of them are verified by dRNA-Seq (**Figure 4**). These splice junctions are used by at least twenty HSV-1 transcripts. Approximately a hundred additional, potential splices sites were identified by the LoRTIA package (**Supplementary Table 1**), which were not confirmed by dRNA-Seq. However, dRNA-Seq produced relative low read coverage, which probably account for this failure. By far the most complex splicing pattern was detected in RNAs produced from the *ul52-54* genomic region.

**Table 6.** The most frequent splice sites of the HSV-1 transcriptome. The newly discovered splice sites are labeled by asterisks.

| Intron start | Intron end | Read count | Splice donor/acceptor | dRNA-Seq |  |
| --- | --- | --- | --- | --- | --- |
| 2318 | 3082 | 20 | GT/AG | dRNA-Seq |  |
| 113428 | 113786 | 40 | GT/AG | dRNA-Seq | * |
| 111360 | 113786 | 9 | GT/AG | - |  |
| 109131 | 111080 | 7 | GT/AG | - | * |
| 97724 | 97949 | 228 | GT/AG | dRNA-Seq |  |
| 91553 | 92535 | 120 | GT/AG | - |  |
| 145649 | 145820 | 1077 | GT/AG | - |  |
| 132373 | 132509 | 2802 | GT/AG | - |  |
| 91123 | 91413 | 95 | GT/AG | - |  |
| 30049 | 33634 | 198 | GT/AG | dRNA-Seq |  |
| 3750 | 3885 | 8 | GT/AG | dRNA-Seq |  |
| 69670 | 69923 | 20 | GT/AG | dRNA-Seq | * |
| 69593 | 69923 | 12 | GT/AG | dRNA-Seq | * |

**Figure 4.**
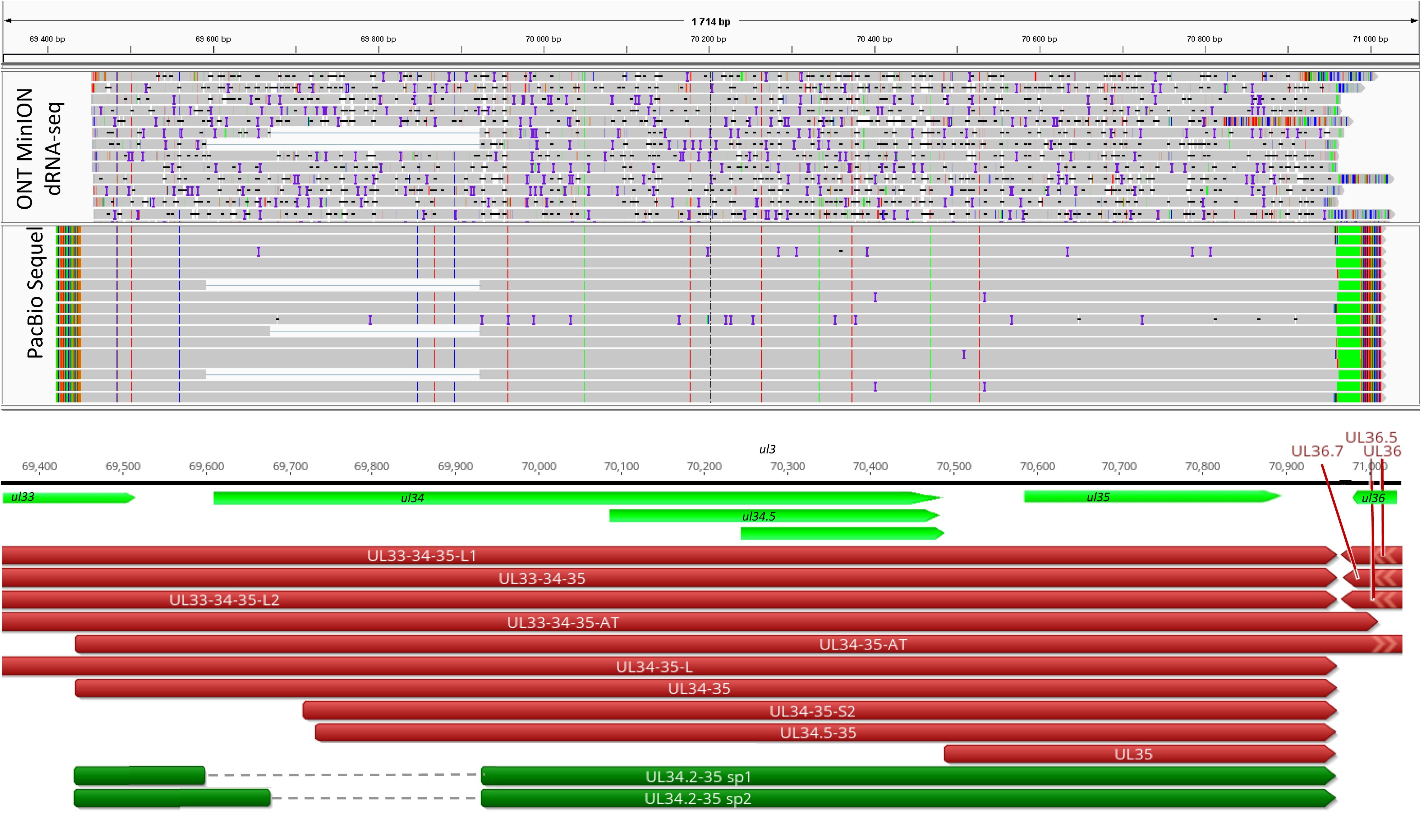
Splice sites of the UL34-35 transcript confirmed by dRNA. Color legend: light green - ORF; red - mRNA; dark green – novel spliced transcripts

### Novel multigenic transcripts

Our earlier survey has revealed several multigenic RNAs, including polycistronic and complex transcripts (Tombácz et al., 2017b). In this work, we identified 126 multigenic transcripts containing two or more genes (**Supplementary Table 5**). The cxRNAs are long RNA molecules with at least two genes standing in opposite orientation relative to one another. An intriguing finding is the RL1-RL2 (ICP34,5-ICP0) bicistronic transcript, as well as the 0.7 kb LAT-UL1-2 and 0.7. kb LAT-UL1-2-3-3.5 cxRNAs (**Figure 1**). Most of the novel multigenic transcripts are expressed at low levels, which can explain why they had previously gone undetected. In this work, we also identified four novel complex transcripts (0.7 kb LAT-UL1-2-c, UL18-15.5-c, UL20-21-c, US4-3-2-c) with unannotated TSSs (**Figure1** and **Supplementary Table 5**). Our novel experiments validated the previously published cxRNAs (**Supplementary Table 5**). This study demonstrates that full-length overlaps between two divergently-oriented HSV-1 genes are an important source for the cxRNA molecules. The likely reason for the lack of cxRNA TSSs is that they are very long and low-abundance transcripts. It cannot be excluded with absolute certainty that some of the low-abundance multigenic transcripts are artefacts produced by the template-switch mechanism; other approaches are needed for the validation of their existence one-by-one.

### Novel transcriptional overlaps

This study revealed an immense complexity of transcriptional overlaps (**Table 7**). These overlaps are produced by either transcriptional read-through between transcripts oriented in parallel or convergent manners (thereby generating rtRNAs), or through using long TSS isoforms of one or of both partners of divergently-oriented genes. Besides these ‘soft’ (alternative) overlaps, adjacent genes can also produce ‘hard’ overlaps when no non-overlapping transcripts are produced from the same gene pairs. An important novelty of this study is the discovery that practically each convergent gene produces rtRNAs crossing the boundaries of the adjacent genes. Two of the convergent gene pairs (*ul3*-*ul4* and *ul30*-*ul3*1) form ‘hard’ transcriptional overlaps, whereas the other gene pairs form ‘soft’ overlaps. The ‘softly’ overlapping convergent transcripts are likely non-polyadenylated, since we were able to detect them by random primer-based sequencing technique alone. The *ul3-ul4* and *ul30-31* gene pairs also express non-polyadenylated rtRNAs that extend beyond their poly(A) sites. Transcriptional read-troughs were detected between each convergent gene pair in most of the cases from both directions, except the UL43-44-45/UL48-47-46 cluster (**Table 7**). Another important novelty of this study is the discovery of very long TSS variants of the divergent transcripts, of which 5’-UTRs entirely overlap the partner gene. We have detected very long transcripts, which overlap the following divergent gene clusters: *ul4-5/ul6-7, ul4-5/ul6-7, ul4-5/ul6-7, ul4-5/ul6-7, ul9-8/ul10, ul9-8/ul10, ul14-13-12-11/ul15, ul17/ul15e2, ul20-19-18/ul21, ul20-19-18/ul21, ulL23-22/ul24-25-26, ul29/*OriL*/ul30, ul29/*OriL*/ul30, ul32-31/ul33-34-35, ul37/ul38-39-40, ul41-ul42, ul49.5.49/ul50, ul51/ul52-53-54, us2/US3, us2/us3, us2/us3*. Altogether, our results show that practically every nucleotide of the double-stranded HSV-1 DNA is transcribed.

**Table 7.**
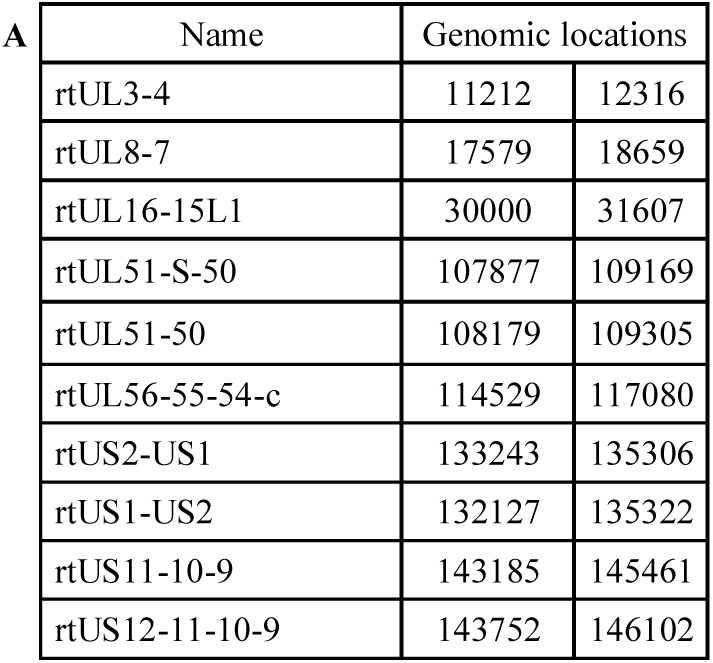
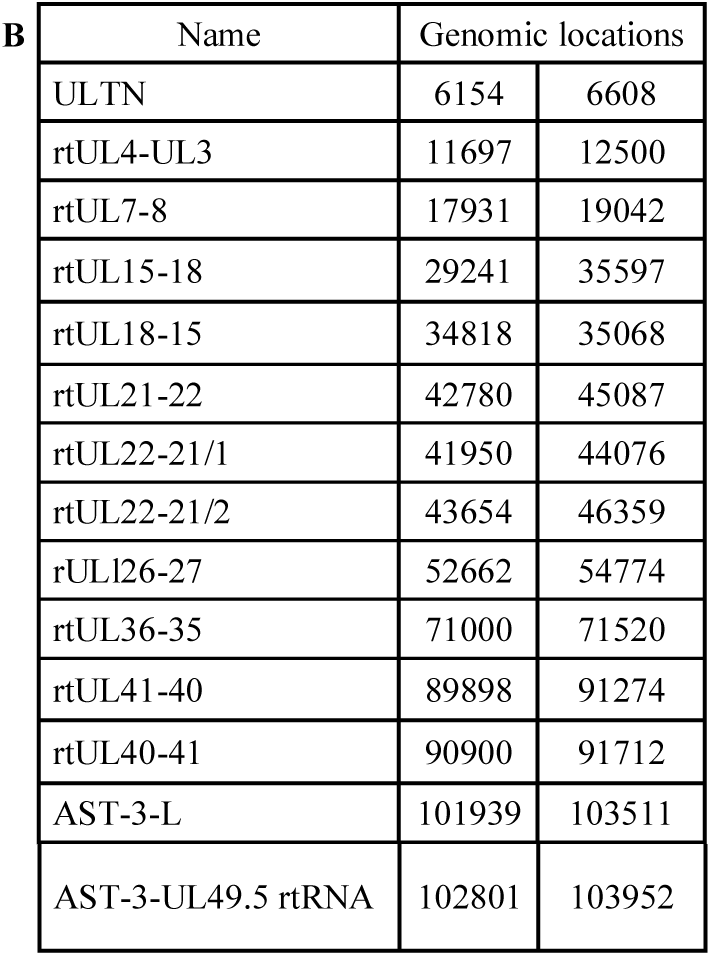
Novel read-through RNAs. **A**: Novel ncRNAs with unidentified 3’ ends; **B**: Novel ncRNAs with unidentified 5’ and 3’ ends. These rtRNAs are probably non-polyadenylated because they were detected by random-primed sequencing alone. The genomic locations indicate the mapping of the transcription reads and not the transcript termini. “rt” stands for “read-through”, “c” for “complex”.

## DISCUSSION

Short-read sequencing has emerged as a golden standard in genomics in recent years (Djebali et al., 2012; Mortazavi et al., 2008; Wang et al., 2009). However, SRS technologies have inherent limitations both in genome and transcriptome analyses. This approach does not perform very well in mapping repetitive elements and GC-rich DNA sequences, or in discriminating paralogous sequences. In transcriptome research, SRS techniques have difficulties in identifying multi-spliced transcripts, overlapping transcripts, TSS and TES isoforms, as well as multigenic RNA molecules. Long-read sequencing can resolve these obstacles. In the last couple of years, LRS approaches have been applied for the analysis of viral transcriptomes, and have obtained a much higher complexity than it had formerly been captured (Tombácz et al., 2018a).

In this study, two sequencing platforms (PacBio Sequel and ONT MinION) were applied for the investigation of the HSV-1 lytic transcriptome, including both poly(A)^+^ and poly(A)^-^ RNAs. This research yielded a number of novel transcripts and transcript isoforms. We identified novel tmRNAs embedded into larger host viral genes. All of these short novel transcripts contain in-frame ORFs, but it does not necessarily mean that they have coding potency. Indeed, most of the tmRNAs are expressed in low abundance, which raises doubts that they code for proteins.

Moreover, we identified two (OriL RNA-1 and OriS RNA-3) novel replication-associated transcripts overlapping OriL and OriS, respectively. Both raRNAs are long TSS isoform produced the neighboring genes, *us1* for OriS, and *ul30* for OriL. Similar transcripts have also been recently described in other viruses (Moldován et al., 2018b; Prazsák et al., 2018; Tombácz et al., 2015, 2018c). Intriguingly, since the replication origin is located at different genomic regions, the sequences of raRNAs are non-homologous within the subfamily of Alphaherpesvirinae. The function of these transcripts may be the regulation of the initiation replication fork as in the bacterial plasmids (Masukata and Tomizawa, 1986; Tomizawa et al., 1981), or the regulation of the orientation of replication through a collision-based mechanism, as suggested in PRV (Boldogkői et al., 2019; Tombácz et al., 2015). In the latter case the raRNAs are mere byproducts of the regulatory mechanism, but it does not exclude that these transcripts have their own function.

This study also identified a large number of transcript length isoforms varying in their TSS or TES, as well as several splice sites and splice isoforms. Here, we also report the identification of several multigenic transcripts including polycistronic and complex transcripts. The existence of cxRNAs, expressed from convergent gene pairs, indicates that transcription does not stop at gene boundaries but occasionally continues across genes standing in opposite directions of one another. We have also detected pervasive antisense transcript expression throughout the entire viral genome especially with the random primer-based sequencing method. The novel antisense RNAs are typically transcriptional read-through products specified by the promoter of neighboring convergent genes. These normally low-abundance non-polyadenylated transcription reads contain varying 3’ends. We also detected novel divergent transcriptional overlaps: in two cases these transcripts appear to be initiated from the 3’-ends of the adjacent genes.

Altogether, this study revealed a much more complex network of transcriptional overlaps than it has been known before. This can lead us to raise the following question: what could be the function of these overlapping RNA molecules? We propose that they are the byproducts of a genome-wide regulatory mechanism based on transcriptional interference between the adjacent and distal genes as described earlier (Boldogköi, 2012).

## ACCESSION NUMBER

The PacBio RSII sequencing files and data files have been uploaded to the NCBI GEO repository and can be found with GenBank accession number GSE97785. Data (aligned files) from MinION and Sequel sequencing have been deposited to the European Nucleotide Archive.

## ACKNOWLEDGEMENTS

We would like to thank Marianna Ábrahám (University of Szeged) for technical assistance. This study was supported by OTKA K 128247 to ZBo and OTKA FK 128252 to DT. DT was also supported by the Bolyai János Scholarship of the Hungarian Academy of Sciences and by the Eötvös Scholarship of the Hungarian State. ZBa was supported by the UNKP-18-3 New National Excellence Program of the ministry of Human Capacities. The project was also supported by the NIH Centers of Excellence in Genomic Science (CEGS) Center for Personal Dynamic Regulomes [5P50HG00773502] to MS.

## AUTHOR CONTRIBUTIONS

DT designed the experiments, prepared the PacBio and ONT sequencing libraries, performed the PacBio RSII, Sequel and ONT MinION sequencing, analyzed the data and drafted the manuscript. ZBa adapted the LoRTIA pipeline for the analysis. GG analyzed the PacBio and ONT dataset and maintained the cell culture. MB analyzed the PacBio data, cultured the Vero cells and made manuscript revisions. ZC isolated the RNAs, generated cDNAs, prepared ONT libraries and performed ONT MinION sequencing. MS conceived and designed the experiments. ZBo conceived and designed the experiments, supervised the study, analyzed the data and wrote the final manuscript. All authors have read and approved the final version of the manuscript.

## SUPPLEMENTARY MATERIAL

**Supplementary Table 1. Summary table of the potential splice junctions of HSV-1 transcripts annotated by LoRTIA program package.** The results from the different library preparation methods are available in distinct excel sheets.

**Supplementary Table 2. The annotated transcriptional starts sites of HSV-1 transcripts.** The potential TS positions detected by using the LoRTIA tool from the different samples (libraries) are available in distinct excel sheets.

**Supplementary Table 3. The transcriptional end sites of HSV-1 transcripts annotated by LoRTIA program package.** The results from the applied sequencing methods are labeled by different colors. Black: Sequel; Blue: RSII; Red: MinION; Green: CAP

**Supplementary Table 4.**. **Long and short TSS isoforms of the HSV-1 transcripts**. Isoforms with unannotated 5’-ends are labeled with asterisks. The genomic positions of these latter transcripts indicate the positions of the longest reads obtained in our experiments. The asterisks indicate unidentified TSSs.

**Supplementary Table 5.**. **Multigenic transcripts and transcript isoforms detected by the different long-read sequencing methods.** From the 125 multigenic RNA molecules 43 isoforms are novel (labeled by asterisk). A ‘c’ letter was added to the transcript names in order to indicate that they are cxRNAs. The cxRNAs with unidentified TSS is labeled with exclamation marks. “sp” stands for “spliced”; “L” for “long”; “S” for “short”; “AT” for “alternative termination”; “c” for “complex”; and “rt” for “read-through”. The asterisks indicate the novel transcripts.

## REFERENCES

Balázs, Z., Tombácz, D., Szűcs, A., Csabai, Z., Megyeri, K., Petrov, A. N., et al. (2017a). Long-Read Sequencing of Human Cytomegalovirus Transcriptome Reveals RNA Isoforms Carrying Distinct Coding Potentials. Sci. Rep. 7, 15989. doi 10.1038/s41598-017-16262-z.

Balázs, Z., Tombácz, D., Szűcs, A., Snyder, M., and Boldogkői, Z. (2017b). Long-read sequencing of the human cytomegalovirus transcriptome with the Pacific Biosciences RSII platform. Sci. Data 4, 170194. doi:10.1038/sdata.2017.194.

Boldogköi, Z. (2012). Transcriptional interference networks coordinate the expression of functionally related genes clustered in the same genomic loci. Front. Genet. 3, 122. doi:10.3389/fgene.2012.00122.

Boldogkői, Z., Balázs, Z., Moldován, N., Prazsák, I., and Tombácz, D. (2019). Novel classes of replication-associated transcripts discovered in viruses. RNA Biol., 1–10. doi:10.1080/15476286.2018.1564468.

Boldogkői, Z., Moldován, N., Szűcs, A., and Tombácz, D. (2018a). Transcriptome-wide analysis of a baculovirus using nanopore sequencing. Sci. Data 5, 180276. doi:10.1038/sdata.2018.276.

Boldogkői, Z., Szűcs, A., Balázs, Z., Sharon, D., Snyder, M., and Tombácz, D. (2018b). Transcriptomic study of Herpes simplex virus type-1 using full-length sequencing techniques. Sci. Data 5, 180266. doi:10.1038/sdata.2018.266.

Byrne, A., Beaudin, A. E., Olsen, H. E., Jain, M., Cole, C., Palmer, T., et al. (2017). Nanopore long-read RNAseq reveals widespread transcriptional variation among the surface receptors of individual B cells. Nat. Commun. 8, 16027. doi:10.1038/ncomms16027.

Chen, S.-Y., Deng, F., Jia, X., Li, C., and Lai, S.-J. (2017). A transcriptome atlas of rabbit revealed by PacBio single-molecule long-read sequencing. Sci. Rep. 7, 7648. doi:10.1038/s41598-017-08138-z.

Cheng, B., Furtado, A., and Henry, R. J. (2017). Long-read sequencing of the coffee bean transcriptome reveals the diversity of full-length transcripts. Gigascience 6, 1–13. doi:10.1093/gigascience/gix086.

Costa, R. H., Cohen, G., Eisenberg, R., Long, D., and Wagner, E. (1984). Direct demonstration that the abundant 6-kilobase herpes simplex virus type 1 mRNA mapping between 0.23 and 0.27 map units encodes the major capsid protein VP5. J. Virol. 49, 287–92. Available at: http://www.ncbi.nlm.nih.gov/pubmed/6317894 [Accessed March 17, 2017].

Depledge, D. P., Puthankalam, S. K., Sadaoka, T., Beady, D., Mori, Y., Placantonakis, D., et al. (2018). Native RNA sequencing on nanopore arrays redefines the transcriptional complexity of a viral pathogen. bioRxiv, 373522. doi:10.1101/373522.

Djebali, S., Davis, C. A., Merkel, A., Dobin, A., Lassmann, T., Mortazavi, A., et al. (2012). Landscape of transcription in human cells. Nature 489, 101–8. doi:10.1038/nature11233.

Du, T., Han, Z., Zhou, G., Roizman, B., and Roizman, B. (2015). Patterns of accumulation of miRNAs encoded by herpes simplex virus during productive infection, latency, and on reactivation. Proc. Natl. Acad. Sci. 112, E49–E55. doi:10.1073/pnas.1422657112.

Harkness, J. M., Kader, M., and DeLuca, N. A. (2014). Transcription of the herpes simplex virus 1 genome during productive and quiescent infection of neuronal and nonneuronal cells. J. Virol. 88, 6847–61. doi:10.1128/JVI.00516-14.

Hu, B., Huo, Y., Chen, G., Yang, L., Wu, D., and Zhou, J. (2016). Functional prediction of differentially expressed lncRNAs in HSV-1 infected human foreskin fibroblasts. Virol. J. 13, 137. doi:10.1186/s12985-016-0592-5.

Li, H. (2018). Minimap2: pairwise alignment for nucleotide sequences. Bioinformatics 34, 3094–3100. doi:10.1093/bioinformatics/bty191.

Li, Y., Fang, C., Fu, Y., Hu, A., Li, C., Zou, C., et al. (2018). A survey of transcriptome complexity in Sus scrofa using single-molecule long-read sequencing. DNA Res. 25, 421–437. doi:10.1093/dnares/dsy014.

Lim, F. (2013). HSV-1 as a Model for Emerging Gene Delivery Vehicles. ISRN Virol. 2013, 1–12. doi:10.5402/2013/397243.

Looker, K. J., Magaret, A. S., May, M. T., Turner, K. M. E., Vickerman, P., Gottlieb, S. L., et al. (2015). Global and Regional Estimates of Prevalent and Incident Herpes Simplex Virus Type 1 Infections in 2012. PLoS One 10, e0140765. doi:10.1371/journal.pone.0140765.

Macdonald, S. J., Mostafa, H. H., Morrison, L. A., and Davido, D. J. (2012). Genome sequence of herpes simplex virus 1 strain KOS. J. Virol. 86, 6371–2. doi:10.1128/JVI.00646-12.

Masukata, H., and Tomizawa, J. (1986). Control of primer formation for ColE1 plasmid replication: Conformational change of the primer transcript. Cell 44, 125–136. doi:10.1016/0092-8674(86)90491-5.

McGeoch, D. J., Rixon, F. J., and Davison, A. J. (2006). Topics in herpesvirus genomics and evolution. Virus Res. 117, 90–104. doi:10.1016/j.virusres.2006.01.002.

McKnight, S. L. (1980). The nucleotide sequence and transcript map of the herpes simplex virus thymidine kinase gene. Nucleic Acids Res. 8, 5949–5964. doi:10.1093/nar/8.24.5949.

Merrick, W. C. (2004). Cap-dependent and cap-independent translation in eukaryotic systems. Gene 332, 1–11. doi:10.1016/j.gene.2004.02.051.

Miyamoto, M., Motooka, D., Gotoh, K., Imai, T., Yoshitake, K., Goto, N., et al. (2014). Performance comparison of second-and third-generation sequencers using a bacterial genome with two chromosomes. BMC Genomics 15, 699. doi:10.1186/1471-2164-15-699.

Moldován, N., Balázs, Z., Tombácz, D., Csabai, Z., Szűcs, A., Snyder, M., et al. (2017a). Multi-platform analysis reveals a complex transcriptome architecture of a circovirus. Virus Res. 237, 37–46. doi:10.1016/j.virusres.2017.05.010.

Moldován, N., Szűcs, A., Tombácz, D., Balázs, Z., Csabai, Z., Snyder, M., et al. (2018a). Multi-platform Next-generation Sequencing Identifies Novel RNA Molecules and Transcript Isoforms of the Endogenous Retrovirus Isolated from Cultured Cells. FEMS Microbiol. Lett. doi:10.1093/femsle/fny013.

Moldován, N., Tombácz, D., Szűcs, A., Csabai, Z., Balázs, Z., Kis, E., et al. (2018b). Third-generation Sequencing Reveals Extensive Polycistronism and Transcriptional Overlapping in a Baculovirus. Sci. Rep. 8, 8604. doi:10.1038/s41598-018-26955-8.

Moldován, N., Tombácz, D., Szűcs, A., Csabai, Z., Snyder, M., and Boldogkői, Z. (2017b). Multi-platform Sequencing Approach Reveals a Novel Transcriptome Profile in Pseudorabies Virus. Front. Microbiol. 8, 2708. doi:10.3389/FMICB.2017.02708.

Mortazavi, A., Williams, B. A., McCue, K., Schaeffer, L., and Wold, B. (2008). Mapping and quantifying mammalian transcriptomes by RNA-Seq. Nat. Methods 5, 621–8. doi:10.1038/nmeth.1226.

Naito, J., Mukerjee, R., Mott, K. R., Kang, W., Osorio, N., Fraser, N. W., et al. (2005). Identification of a protein encoded in the herpes simplex virus type 1 latency associated transcript promoter region. Virus Res. 108, 101–110. doi:10.1016/j.virusres.2004.08.011.

Nudelman, G., Frasca, A., Kent, B., Sadler, K. C., Sealfon, S. C., Walsh, M. J., et al. (2018). High resolution annotation of zebrafish transcriptome using long-read sequencing. Genome Res. 28, 1415–1425. doi:10.1101/gr.223586.117.

O’Grady, T., Wang, X., Höner zu Bentrup, K., Baddoo, M., Concha, M., and Flemington, E. K. (2016). Global transcript structure resolution of high gene density genomes through multi-platform data integration. Nucleic Acids Res. 44, e145–e145. doi:10.1093/nar/gkw629.

Oláh, P., Tombácz, D., Póka, N., Csabai, Z., Prazsák, I., and Boldogkői, Z. (2015). Characterization of pseudorabies virus transcriptome by Illumina sequencing. BMC Microbiol. 15, 130. doi:10.1186/s12866-015-0470-0.

Perng, G.-C., Maguen, B., Jin, L., Mott, K. R., Kurylo, J., BenMohamed, L., et al. (2002). A novel herpes simplex virus type 1 transcript (AL-RNA) antisense to the 5’ end of the latency-associated transcript produces a protein in infected rabbits. J. Virol. 76, 8003–10. doi:10.1128/JVI.76.16.8003-8010.2002.

Prazsák, I., Moldován, N., Balázs, Z., Tombácz, D., Megyeri, K., Szűcs, A., et al. (2018). Long-read sequencing uncovers a complex transcriptome topology in varicella zoster virus. BMC Genomics 19, 873. doi:10.1186/s12864-018-5267-8.

Rajčáni, J., Andrea, V., and Ingeborg, R. (2004). Peculiarities of Herpes Simplex Virus (HSV) Transcription: An overview. Virus Genes 28, 293–310. doi:10.1023/B:VIRU.0000025777.62826.92.

Rixon, F. J., and Clements, J. B. (1982). Detailed structural analysis of two spliced HSV-1 immediate-early mRNAs. Nucleic Acids Res. 10, 2241–2256. doi:10.1093/nar/10.7.2241.

Sedlackova, L., Perkins, K. D., Lengyel, J., Strain, A. K., van Santen, V. L., and Rice, S. a (2008). Herpes simplex virus type 1 ICP27 regulates expression of a variant, secreted form of glycoprotein C by an intron retention mechanism. J. Virol. 82, 7443–55. doi:10.1128/JVI.00388-08.

Stingley, S. W., Ramirez, J. J., Aguilar, S. A., Simmen, K., Sandri-Goldin, R. M., Ghazal, P., et al. (2000). Global analysis of herpes simplex virus type 1 transcription using an oligonucleotide-based DNA microarray. J. Virol. 74, 9916–27. Available at: http://www.ncbi.nlm.nih.gov/pubmed/11024119 [Accessed February 4, 2016].

Tombácz, D., Balázs, Z., Csabai, Z., Moldován, N., Szűcs, A., Sharon, D., et al. (2017a). Characterization of the Dynamic Transcriptome of a Herpesvirus with Long-read Single Molecule Real-Time Sequencing. Sci. Rep. 7, 43751. doi:10.1038/srep43751.

Tombácz, D., Balázs, Z., Csabai, Z., Snyder, M., and Boldogkői, Z. (2018a). Long-Read Sequencing Revealed an Extensive Transcript Complexity in Herpesviruses. Front. Genet. 9, 259. doi:10.3389/fgene.2018.00259.

Tombácz, D., Csabai, Z., Oláh, P., Balázs, Z., Likó, I., Zsigmond, L., et al. (2016). Full-Length Isoform Sequencing Reveals Novel Transcripts and Substantial Transcriptional Overlaps in a Herpesvirus. PLoS One 11, e0162868. doi:10.1371/journal.pone.0162868.

Tombácz, D., Csabai, Z., Oláh, P., Havelda, Z., Sharon, D., Snyder, M., et al. (2015). Characterization of Novel Transcripts in Pseudorabies Virus. Viruses 7, 2727–2744. doi:10.3390/v7052727.

Tombácz, D., Csabai, Z., Szűcs, A., Balázs, Z., Moldován, N., Sharon, D., et al. (2017b). Long-Read Isoform Sequencing Reveals a Hidden Complexity of the Transcriptional Landscape of Herpes Simplex Virus Type 1. Front. Microbiol. 8, 1079. doi:10.3389/fmicb.2017.01079.

Tombácz, D., Prazsák, I., Moldován, N., Szűcs, A., and Boldogkői, Z. (2018b). Lytic Transcriptome Dataset of Varicella Zoster Virus Generated by Long-Read Sequencing. Front. Genet. 9, 460. doi:10.3389/fgene.2018.00460.

Tombácz, D., Prazsák, I., Szűcs, A., Dénes, B., Snyder, M., and Boldogkői, Z. (2018c). Dynamic transcriptome profiling dataset of vaccinia virus obtained from long-read sequencing techniques. Gigascience 7. doi:10.1093/gigascience/giy139.

Tombácz, D., Sharon, D., Szűcs, A., Moldován, N., Snyder, M., and Boldogkői, Z. (2018d). Transcriptome-wide survey of pseudorabies virus using next-and third-generation sequencing platforms. Sci. Data 5, 180119. doi:10.1038/sdata.2018.119.

Tombácz, D., Tóth, J. S., Petrovszki, P., and Boldogkői, Z. (2009). Whole-genome analysis of pseudorabies virus gene expression by real-time quantitative RT-PCR assay. BMC Genomics 10, 491. doi:10.1186/1471-2164-10-491.

Tomizawa, J., Itoh, T., Selzer, G., and Som, T. (1981). Inhibition of ColE1 RNA primer formation by a plasmid-specified small RNA. Proc. Natl. Acad. Sci. U. S. A. 78, 1421–5. doi:10.1073/PNAS.78.3.1421.

Voss, J. H., and Roizman, B. (1988). Properties of two 5’-coterminal RNAs transcribed part way and across the S component origin of DNA synthesis of the herpes simplex virus 1 genome. Proc. Natl. Acad. Sci. U. S. A. 85, 8454–8. Available at: http://www.ncbi.nlm.nih.gov/pubmed/2847162 [Accessed December 18, 2016].

Wang, Z., Gerstein, M., and Snyder, M. (2009). RNA-Seq: a revolutionary tool for transcriptomics. Nat. Rev. Genet. 10, 57–63. doi:10.1038/nrg2484.

Wen, M., Ng, J. H. J., Zhu, F., Chionh, Y. T., Chia, W. N., Mendenhall, I. H., et al. (2018). Exploring the genome and transcriptome of the cave nectar bat Eonycteris spelaea with PacBio long-read sequencing. Gigascience 7. doi:10.1093/gigascience/giy116.

Zhang, B., Liu, J., Wang, X., and Wei, Z. (2018). Full-length RNA sequencing reveals unique transcriptome composition in bermudagrass. Plant Physiol. Biochem. 132, 95–103. doi:10.1016/j.plaphy.2018.08.039.

Zhu, Y. Y., Machleder, E. M., Chenchik, A., Li, R., and Siebert, P. D. (2001). Reverse transcriptase template switching: a SMART approach for full-length cDNA library construction. Biotechniques 30, 892–7. Available at: http://www.ncbi.nlm.nih.gov/pubmed/11314272.

